# Predicting functional consequences of mutations using molecular interaction network features

**DOI:** 10.1101/2021.03.05.433991

**Authors:** Kivilcim Ozturk, Hannah Carter

## Abstract

Variant interpretation remains a central challenge for precision medicine. Missense variants are particularly difficult to understand as they change only a single amino acid in protein sequence yet can have large and varied effects on protein activity. Numerous tools have been developed to identify missense variants with putative disease consequences from protein sequence and structure. However, biological function arises through higher order interactions among proteins and molecules within cells. We therefore sought to capture information about the potential of missense mutations to perturb protein interaction networks by integrating protein structure and interaction data. We developed 16 network-based annotations for missense mutations that provide orthogonal information to features classically used to prioritize variants. We then evaluated them in the context of a proven machine-learning framework for variant effect prediction across multiple benchmark datasets to demonstrate their potential to improve variant classification. Interestingly, network features resulted in larger performance gains for classifying somatic mutations than for germline variants, possibly due to different constraints on what mutations are tolerated at the cellular versus organismal level. Our results suggest that modeling variant potential to perturb context-specific interactome networks is a fruitful strategy to advance *in silico* variant effect prediction.

## INTRODUCTION

Advances in high throughput sequencing technologies have resulted in the rapid accumulation of genomic data and allowed profiling of patient genomes in clinical settings. Such studies frequently uncover previously unobserved and uncharacterized genetic variants of ambiguous relevance to health, making variant interpretation an important challenge in precision medicine [1]. Missense mutations are particularly challenging as they only change a single amino acid in a protein sequence yet can have effects spanning no difference to complete loss of function. Numerous methods have been developed to prioritize functional missense variants [2–10]. Typically, these tools rely on protein sequence/structure information to predict variant effects at the protein level, and the scores they provide tend to capture coarse grained estimates of impact (e.g damaging, benign, tolerated).

Biological functions and cellular behaviors arise from interactions among proteins and other molecules within cells, and biological systems evolve to be robust to random error [11]. Diseases are often associated with perturbations to protein interactions, different perturbations can result in different phenotypes [12], and the level of impact caused by mutations to the underlying molecular interaction network may determine the likelihood of generating a phenotype [13]. For example, loss of function mutations were more likely to be tolerated when they affected proteins at the periphery of the interactome [14]. Similarly, variants that otherwise were predicted to have little effect were more likely to be deleterious if they had a large number of interaction partners [15]. Thus, a protein’s location within the system provides biological context that may be important for understanding the effects of mutations [16].

Within proteins, different mutations may have different effects on protein functions [17,18]. While destabilizing mutations at the core of a protein are likely to interfere with all protein activities, mutations on the surface could potentially interfere with specific protein activities while preserving others [17]. In this way, different mutations targeting the same protein might perturb its interactions differently, affecting different pathways that the protein is involved in, and resulting in different disease phenotypes [19]. Indeed, analyses have demonstrated an unexpected enrichment of Mendelian mutations [20–22], and somatic mutations [23–26] at protein interaction interfaces.

Based on the above we sought to assess the potential for artificial intelligence based methods for variant interpretation to derive new information from molecular interaction data. We first integrated structure and protein-protein interaction (PPI) networks to enable systematic annotation of proteins according to location and interactions (Figure 1A). We mapped various germline variants and somatic mutations to network edges to describe their potential to impact biological function (Figure 1B). We then designed features capturing information about proteins and amino acids in the context of their importance to the network architecture and evaluated them within a machine-learning variant classification framework (Figure 1C). We found that network-based features capture orthogonal information to classical amino acid (AA) sequence/structure-based features and can improve variant classification, though they may be more informative for some variant classification tasks than others.

**Figure 1.**
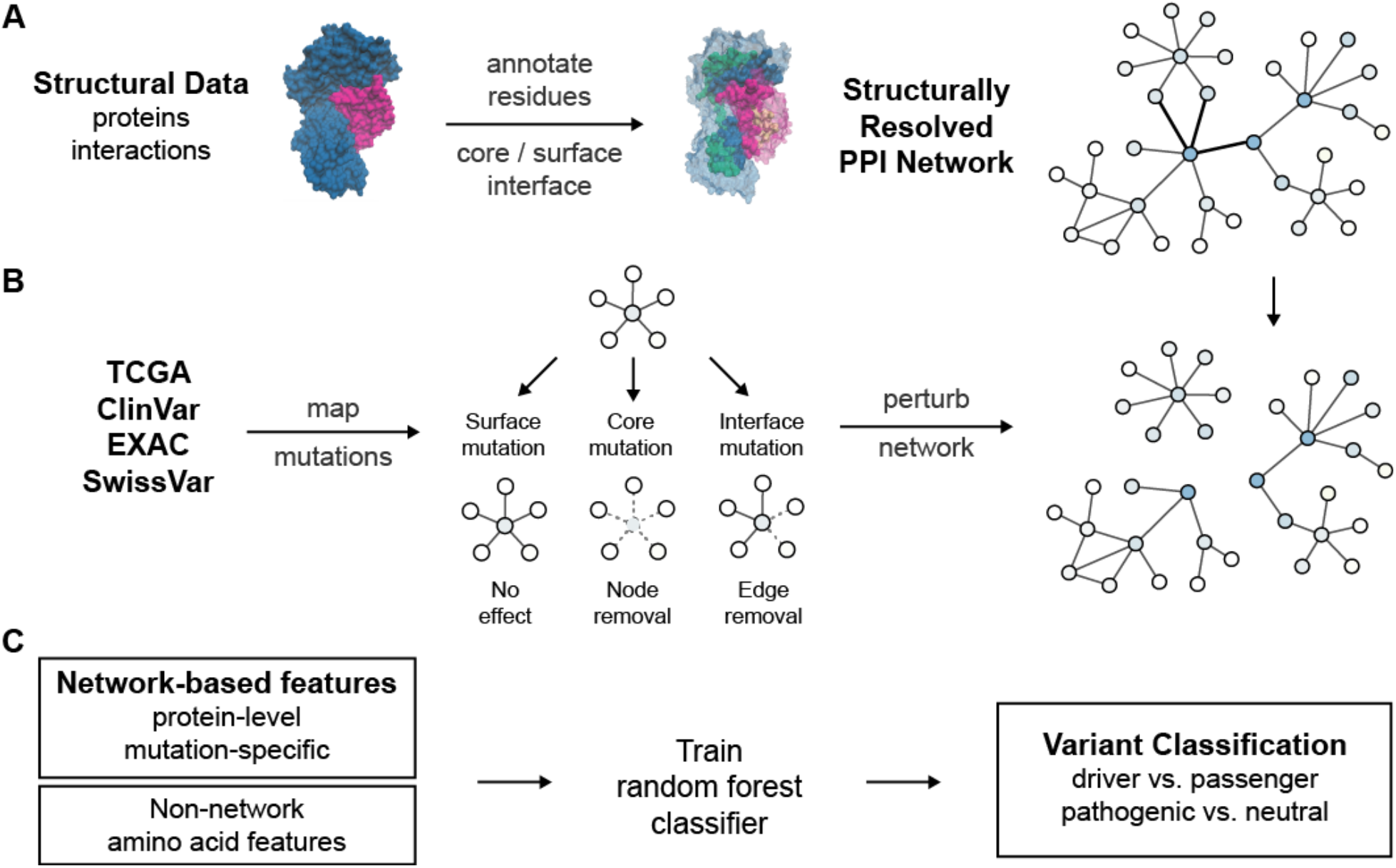
Overview of the method. **(A)** Constructing a structurally resolved PPI network. **(B)** Mapping mutations to perturbed network architectures. **(C)** Designing protein-level and mutation-specific network-based features and using a machine learning framework to evaluate their potential for variant classification alone and in combination with classic non-network amino acid features.

## RESULTS

### Disease causing genes are central in PPI networks

The architectures of biological networks can provide important information for understanding the pathogenesis of mutations [16,27]. The scale-free topology of PPI networks suggests that they are more tolerant to random failures, but variants affecting higher degree nodes are more likely to disrupt function [28]. Indeed, when we compared disease genes using a high-confidence human PPI network of experimentally verified interactions from STRING [29], cancer driver [30] and Mendelian disease genes [31] score higher with various centrality measures than other genes (Figure 2). This suggests that the network niche of a gene provides information about the potential of an amino acid substitution to create deleterious phenotypes, a relationship that has proven robust to study bias [32]; in our data only node degree correlates weakly with the number of publications (Pearson r=0.23, Figure S1).

**Figure 2.**
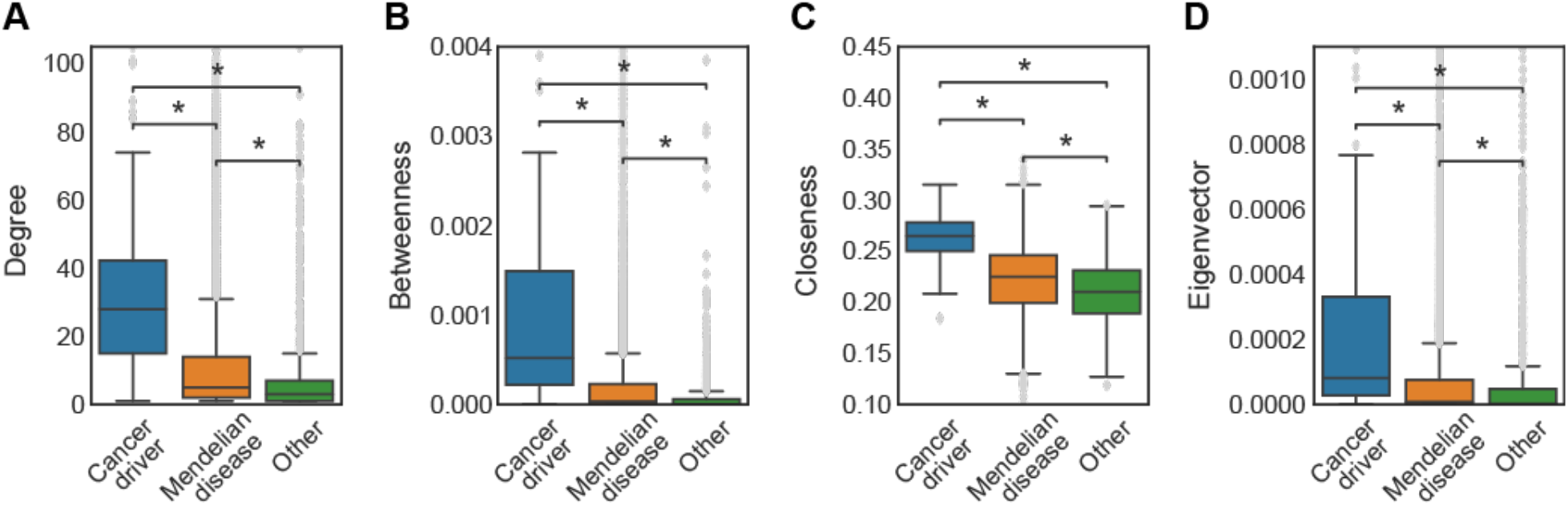
Disease genes are central in PPI networks. Boxplots showing distributions of **(A)** degree, **(B)** betweenness, **(C)** closeness, and **(D)** eigenvector centralities of cancer driver, Mendelian disease and other genes (Mann-Whitney U test; *p<1e-04).

### Creating a structurally resolved PPI network

While disease mutations target proteins more central in interaction networks (Figure 2), protein level descriptors of centrality are not capable of distinguishing the effects of different mutations within proteins. Investigation of mutation-specific network perturbations requires mapping of mutations to 3D protein structures and interaction interfaces so that we can model their potential to affect network edges (Figure 1B). We constructed a structurally resolved PPI network (called SRNet from here on) comprising 6,230 proteins and 10,615 PPIs using 3D structures and homology models (Figure 1A). This network contains annotations for 530,668 interface residues, defined here as the subset of amino acid residues that mediate physical contact between proteins. Otherwise, amino acids are annotated according to location at surface or core based on relative solvent accessible surface area calculated from protein 3D structures (Methods). SRNet is an updated and extended version of our previous structurally resolved PPI network [23].

### Disease mutations frequently target interface or core residues

We further assessed the potential for SRNet to capture information about mutation-based network-perturbation by analyzing location of mutations relative to core, surface or interface regions. Similar to the finding in Engin *et al*. [23], SRNet supports that somatic missense mutations in tumors (obtained from The Cancer Genome Atlas (TCGA) [33]) target surface regions in oncogenes (OR=1.32, p=1.4e-06) and other genes (OR=1.15, p=1.07e-59), but are relatively depleted at surface regions in tumor suppressor genes (OR=0.91, p<0.1) due to a larger proportion of core mutations (Figure 3A), consistent with more loss of function mutations in tumor suppressors. However, when focusing only on surface positions, somatic mutations are more likely to be found at interface regions of oncogenes (OR=1.11, p<0.05) and tumor suppressors (OR=1.30, p=7.8e-07) relative to other genes (Figure 3B). Analysis of pathogenic germline variants (ClinVar [34]) versus neutral variants (EXAC [35], SwissVar [36], ClinVar [34]) found similar trends. Pathogenic variants were relatively depleted at the surface (OR=0.56, p=1.5e-42), suggesting they were far more likely to affect core regions, whereas neutral variants were biased toward the surface (OR=1.69, p=1e-19) (Figure 3C). On protein surfaces, pathogenic variants were more often found at interface regions (OR=5.65, p=2.2e-308) though neutral variants also showed increased odds of affecting an interface (OR=2.87, p=2.6e-115) (Figure 3D).

**Figure 3.**
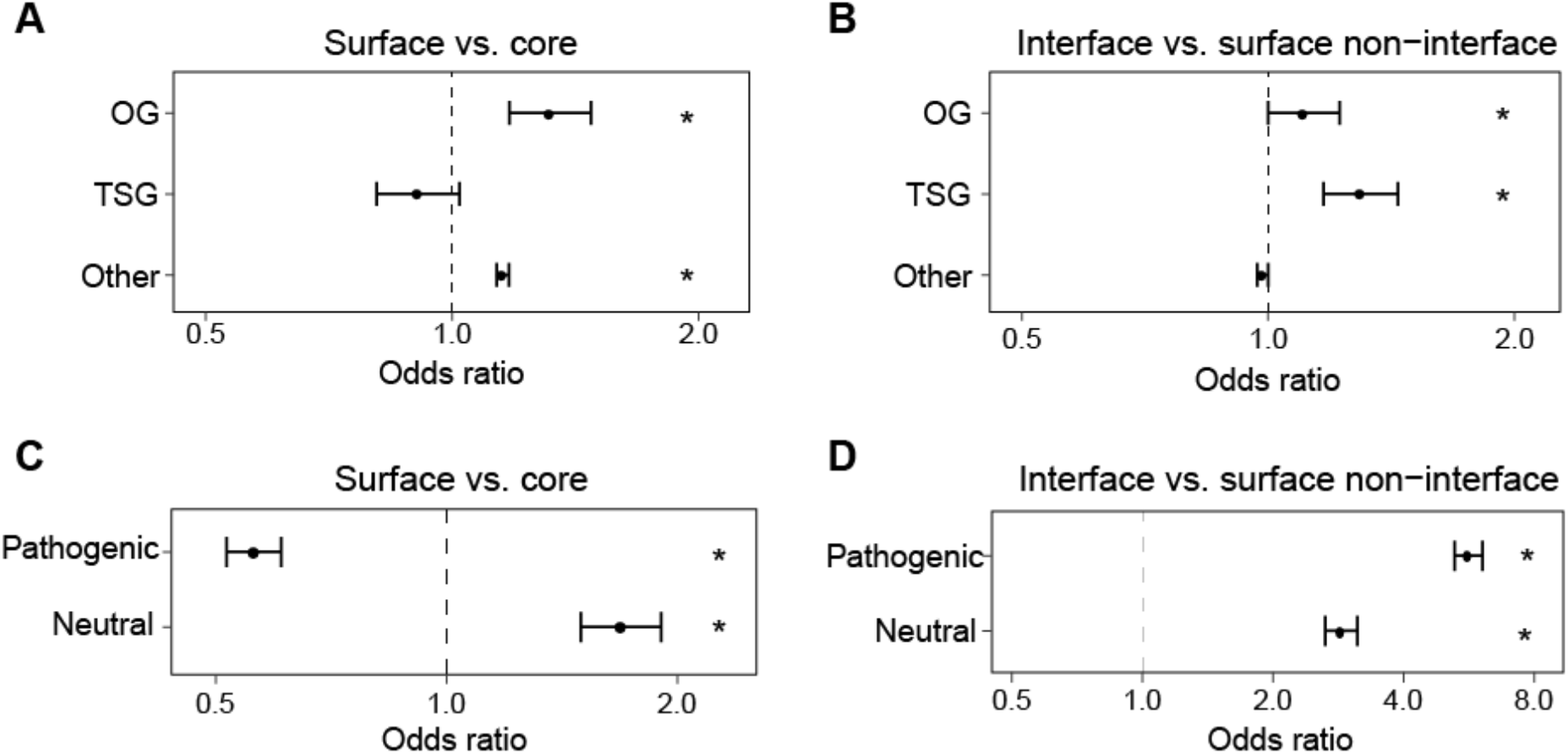
Analysis of structural location of missense disease mutations. Odds ratios (OR) and 95% confidence intervals using Fisher’s exact test are shown (*p<0.05). **(A-B)** Comparison of somatic mutations in oncogenes (OG), tumor suppressor genes (TSG), and other genes located at **(A)** surface vs. core residues, **(B)** surface interface vs. surface non-interface residues. **(C-D)** Comparison of pathogenic and neutral variants located at **(C)** surface vs. core residues, **(D)** surface interface vs. surface non-interface residues. For **(A)** and **(C)**, an OR>1 means more mutations/variants were found at the surface. For **(B)** and **(D)** an OR>1 means more mutations/variants were found at interfaces.

### Network-based features for variant classification

As the above analyses support that both protein and amino acid level information derived from networks is informative about disease-association, we hypothesized that network information would be useful for machine-learning-based variant classification. We designed and analyzed 16 features describing network-level effects of mutations, including 7 protein-level features (Figure 4A) that estimate the significance of the target protein in the network, and 9 mutation-level features (Figures 4B-C) quantifying the potential of individual amino acid positions on the protein to impact network architecture. The mutation-level features are based on comparing network measures before and after removing edges in SRNet potentially affected by a mutation. These 16 features show potential to distinguish between different classes of variants (Figures 4A-C) and are not strongly correlated with other classic non-network based amino acid features used for variant classification, such as measures of site-specific conservation (Figure 4D), suggesting that they add new and useful information.

**Figure 4.**
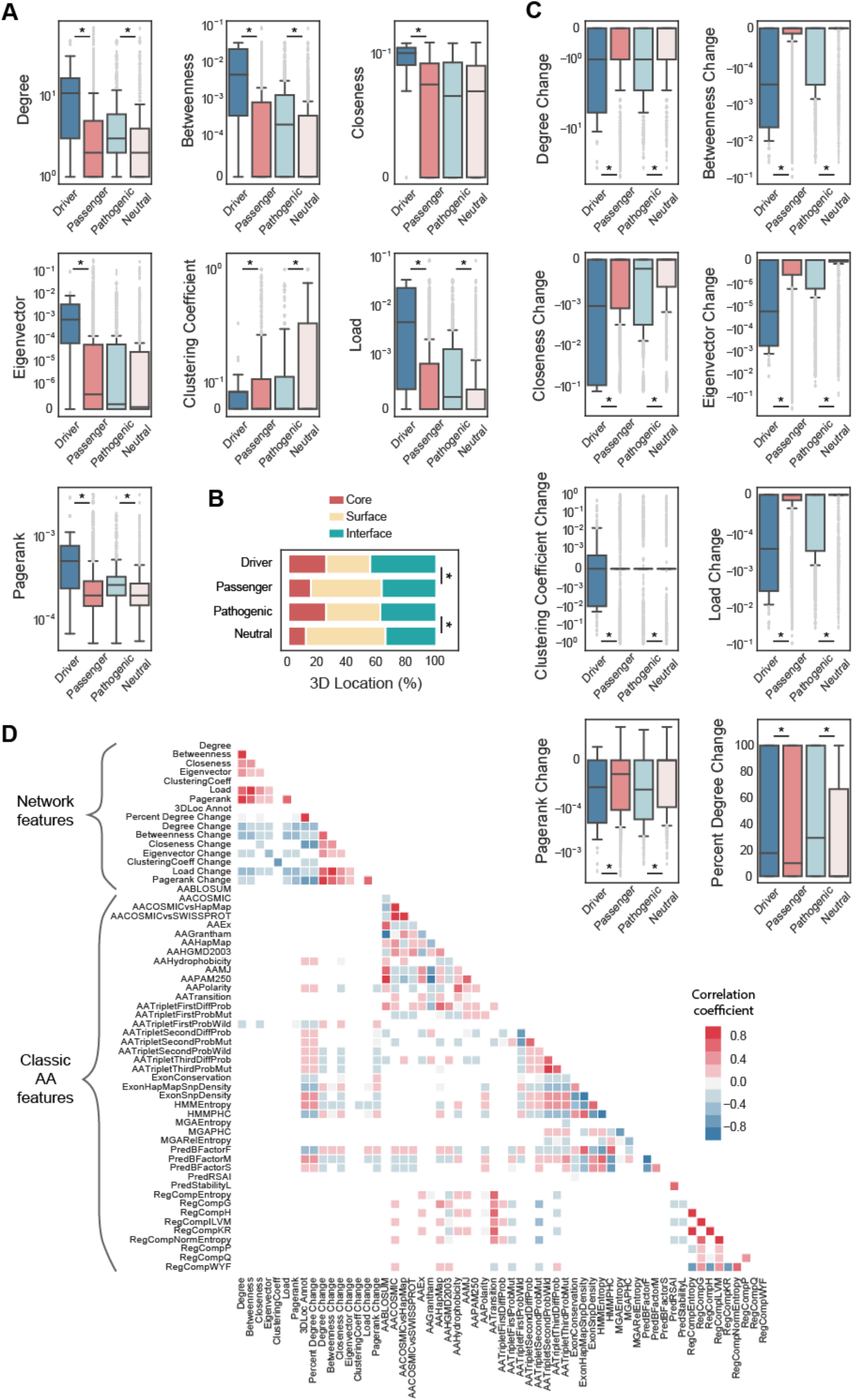
Distribution of network-based features. Distribution of network-based features for driver vs. passenger mutations and pathogenic vs. neutral variants in SRNet (Mann-Whitney U test; *p<1e-04). **(A)** Boxplots showing distribution of protein-level features (degree, betweenness, closeness, eigenvector, clustering coefficient, load and pagerank), **(B)** a stacked bar plot showing percent distribution of 3D locations (core, interface, surface), and **(C)** boxplots showing distribution of mutation-level network features (degree change, betweenness change, closeness change, eigenvector change, clustering coefficient change, load change, pagerank change and percent degree change). **(D)** Heatmap displaying Pearson correlation coefficients of network-based and classic non-network based amino acid features. Only features that have at least one correlation coefficient higher than 0.3 are shown.

### Utility of network features for classifying cancer driver mutations

To evaluate the benefit of using network features for somatic mutation classification, we trained a Random Forest to predict driver or passenger class labels using different combinations of features. We separately evaluated classifier performance when trained using all 16 network derived features, only the 7 protein-level features or the 9 mutation-level features alone, or in combination with 83 amino acid level features obtained from the SNVBox database [37]. As a training set, we used likely-driver and likely-passenger missense mutations from Tokheim *et al*., which they obtained from TCGA using a semi-supervised approach based on known cancer driver gene annotations and mutation rates with the goal of generating a more balanced training set consisting of both driver and passenger mutations in cancer genes [38]. Generalization error was estimated using a 5-fold cross-validation with gene hold out to prevent information leakage and consequent overfitting. We measured performance using the area under the ROC curve (auROC) metric, similar to prior variant effect prediction studies [8]. We note that use of network features limits training and prediction to mutations that can be annotated by SRNet.

For driver classification, protein-level network features performed better than mutation-specific features (Figure 5A), though performance for mutation-level features was better for the top scoring ∼20% of drivers (left edge of ROC curve). We note that mutation-level features alone classify all surface non-interface mutations as passengers since their feature values should all be the same (there is no change to network centrality measures when no edges are affected). Combining mutation-level features with more classic amino acid level features significantly boosts performance over mutation-level features alone (Figure 5B). Interestingly, network features alone slightly outperform amino acid level features alone, pointing to the extreme centrality of driver genes. As mutation-level features are likely to be most informative for mutations at interfaces, we further explored performance for interface mutations only (Figure 5C). Here we see that mutation-specific network features perform considerably better as they are not hindered by misclassification of surface mutations (Figure 5C). Overall, the combination of network-based and amino acid features displays the highest performance (Figures 5B-C).

**Figure 5.**
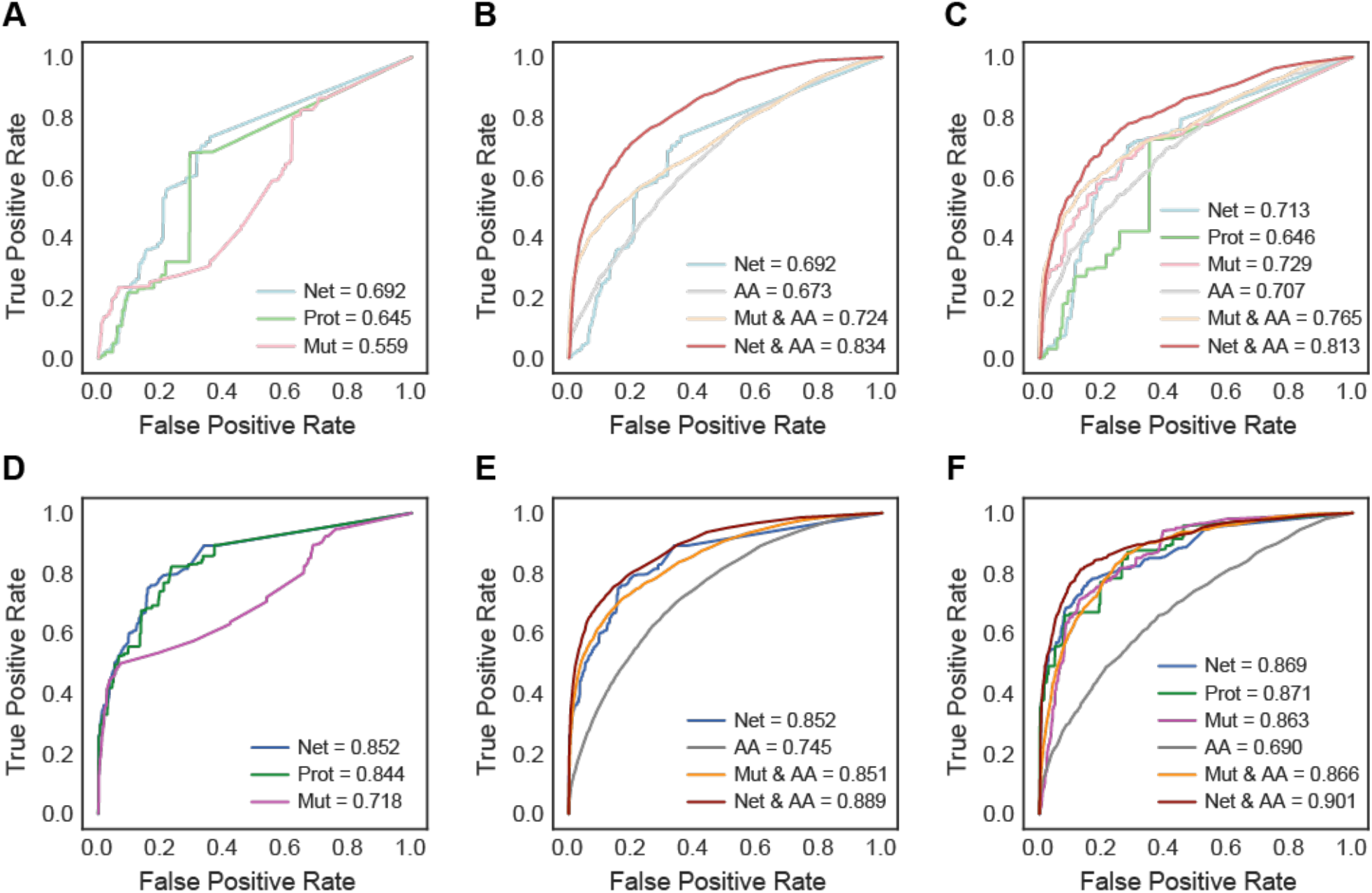
Classifier performance in identifying cancer mutations. ROC curves for identifying cancer mutations using **(A-B-C)** SRNet, and **(D-E-F)** the extended network, with **(A-D)** network (Net), protein-level (Prot) and mutation-level (Mut) features; with **(B-E)** network, amino acid (AA), mutation-level and amino acid (Mut & AA), and network and amino acid (Net & AA) features. **(C-F)** ROC curves for identifying cancer mutations targeting interface residues only using all mentioned features.

### Incorporating in silico predicted interface residues

The restriction to analysis of mutations for which 3D structural information about interfaces is available is a problematic limitation. *In silico* prediction of interfaces can be used to augment interface coverage, such as was done for Interactome INSIDER [39]. To explore whether *in silico* predicted interfaces could boost mutation coverage without loss of performance, we repeated our analysis on an extended network with both structure-derived and predicted interfaces. This resulted in improved performance overall (Figures 5D-F), suggesting that improvements to interface features and the ability to train on a larger set of mutations enabled by higher coverage in the expanded network outweigh the introduction of noise caused by interface prediction error. A more stringent comparison considering only proteins shared between the original and the expanded network found similar results (auROC is 0.832 and 0.871 for SRNet and the extended network for the classifier with Network & AA features respectively).

We further investigated mutation-level network features in the extended network by examining cases where the classifier was successful in differentiating between driver and passenger mutations occurring in the same proteins. Focusing on 150 interface mutations from 8 cancer genes (EGFR, HRAS, KRAS, TP53, RB1, PIK3R1, CTNNB1, EP300) with correct classification of both driver and passenger labels, we observed a significant difference in distribution of mutation-level features for these two classes (Figure 6), further supporting that mutation-level network features provide information useful for within gene mutation classification.

**Figure 6.**
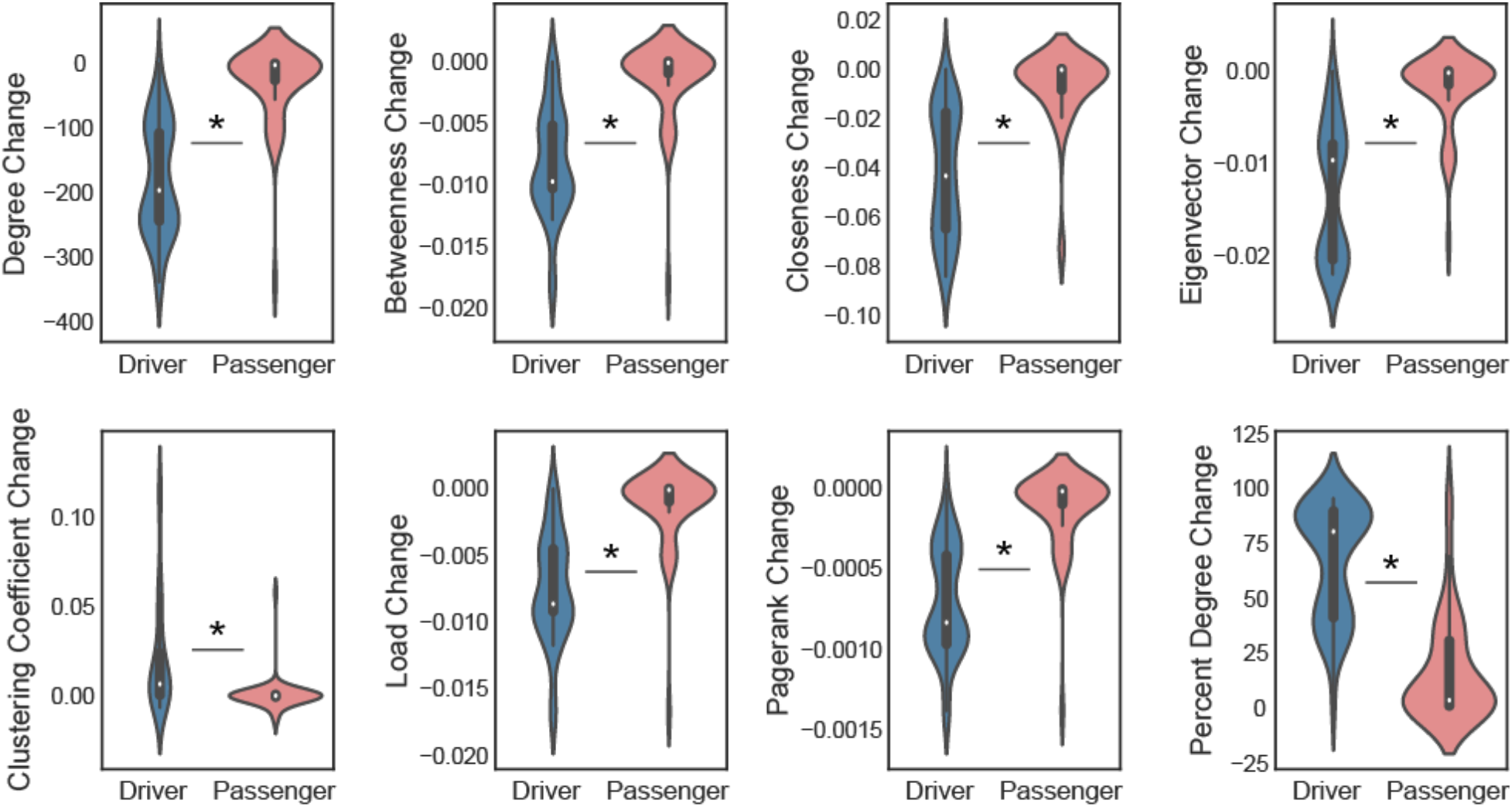
Distribution of mutation-level network features. Distribution of mutation-level network features for correctly labelled driver and passenger mutations occurring in the same proteins (Mann-Whitney U test; *p<1e-04).

### Overall performance on benchmark datasets

We next sought to evaluate the improvement obtained from network features on independent studies of cancer mutations. Since no ‘gold standard’ dataset exists for cancer, we evaluated classifier performance relative to best-in-class methods that do not use network-derived features on 4 external pan-cancer datasets constructed using different approaches: an *in vivo* screen: Kim *et al*. [40], an *in vitro* assay: Ng *et al*. [41], and 2 literature-derived datasets: MSK-IMPACT and CGC-recurrent, previously described in Tokheim *et al*. [38]. For each dataset, classifier performance was evaluated using the area under the ROC (auROC) and PR curves (auPRC).

Using the classifier trained on the extended network described in the previous section, we observed comparable performance to both cancer-specific: CHASM [42], TransFIC [43], FATHMM [44], and CanDrA [45]; and population-based methods: SIFT [4], Polyphen2 [5], VEST [46], and REVEL [8] (Figure 7). Our classifier had the highest auROC and auPRC scores on the *in vitro* set, yielded higher scores than most on the literature curated sets, displayed similar performance on the *in vivo* study, and showed the highest mean performance across datasets. This suggests that network-derived features that capture abstract information about the role of proteins in networks and the potential of mutations to perturb this role, are helpful for driver classification across a variety of settings, though the gains over methods trained on classic amino acid based features are modest. We note that the recently released CHASMplus now includes a feature indicating whether mutations occur at a protein-interaction interface, along with other improvements, and has significantly improved performance over the original method. Location at an interface was reported as the second most informative feature after a feature describing within protein clustering of observed mutations [38].

**Figure 7.**
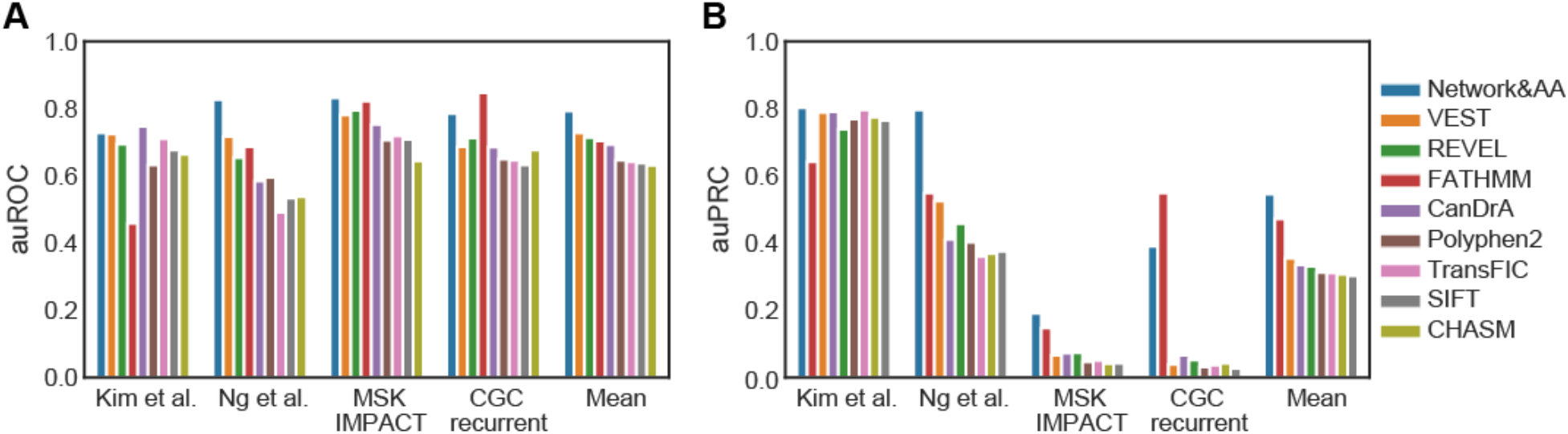
Comparison of classifier performance on benchmark datasets relative to established methods. Bar plots depict the area under the **(A)** ROC (auROC) and **(B)** PR curve (auPRC) scores for each method. Mean category displays the mean of scores of each method across datasets. Classifiers are ordered based on their mean scores. Both panels use the same color scheme displayed on the right.

However, it should also be considered that our network-based approach is dependent on the inclusion of proteins in the network and availability of annotations mapping amino acid residues to core, surface or interface residues. This generally results in a smaller training set than other methods, and an inability to score some fraction of mutations. For the benchmark sets evaluated here, 95.77% of Kim *et al*., 76.57% of Ng *et al*., 70.42% of MSK-IMPACT, and 74.27% of CGC recurrent dataset mutations could be scored respectively. It is possible that better network and amino acid annotation coverage could further boost performance.

### Utility of network features for classifying pathogenic germline variants

We next evaluated whether network features are also useful in the context of germline variation. We previously observed that inherited disease genes were less central than cancer genes and both pathogenic and neutral mutations were enriched at interface positions, though to different extents. We once again trained a Random Forest classifier to prioritize missense mutations that alter protein activity using the 16 network-based features and 83 amino acid descriptors, using a training set composed of pathogenic and neutral variants (described in the section “Disease mutations frequently target interface or core residues”).

For germline variants, mutation-specific features yielded similar (with SRNet) or higher performance (with extended network) than protein-level features for all mutations (Figure 8A), and for interface mutations only (Figure 8C). But overall, network features are outperformed by non-network amino acid features (Figure 8B) which is the opposite of the case with cancer driver classifier. This is consistent with proteins targeted by pathogenic germline variants being less central than cancer driver genes. Since proteins harboring germline pathogenic variants have fewer interaction partners, pathogenic variants in the protein core or at interfaces tend not to result in as extreme values of mutation-level network features as driver mutations do (Figure 4C), despite the observed enrichment for these variants in core and interface regions (Figures 3C-D). The similar performance by network features in both SRNet and the extended network (Figure 8) suggests that either increased coverage does not improve performance as much, or the noise introduced by interface prediction error counteracts the performance gained by higher coverage in this setting. A stricter comparison considering only shared proteins between networks once again showed similar performance (auROC is 0.835 and 0.849 for SRNet and the extended network for the classifier with Network & AA features respectively).

**Figure 8.**
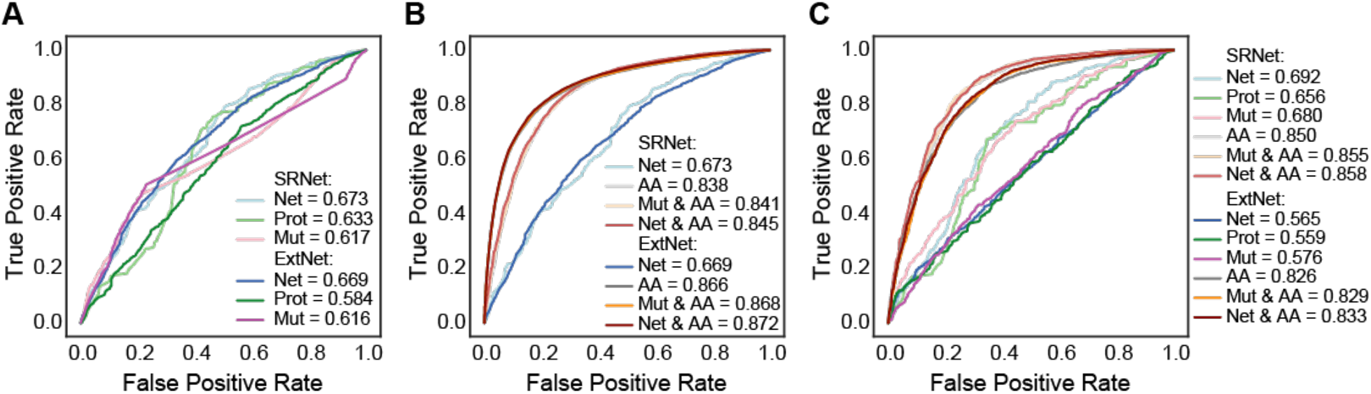
Classifier performances for predicting pathogenic vs. neutral variants using SRNet vs. the extended network (ExtNet). ROC curves for identifying variants with **(A)** network (Net), protein-level (Prot) and mutation-level (Mut) features; with **(B)** network, amino acid (AA), mutation-level and amino acid (Mut & AA), and network and amino acid (Net & AA) features. **(C)** ROC curves for identifying variants targeting interface residues only using all mentioned features.

## DISCUSSION

Understanding the functional consequences of protein coding variants remains a challenging task. The numerous methods developed thus far to predict whether a mutation is likely to impair protein activity or cause a pathogenic phenotype have largely been protein-centric. Systems biology has provided insights as to how molecular interaction networks evolve to ensure robustness to genetic variation [13]. It is increasingly apparent that the role of proteins within molecular networks is a key determinant of the potential of variants to exert deleterious effects [14,15]. Here we show that network-derived features can capture novel information about variant effects that is not already present in the classical amino acid features used by most variant classification methods, and that combining both sets of features improves classifier performance.

Our approach relies on a structurally resolved PPI network that allows variants to be characterized according to their potential to affect network architecture by mapping them to their location on protein structures and protein-interaction interfaces. These mappings are used to capture the potential of variant positions to perturb information flow through the network. We developed protein-level features to capture the relative importance of a protein within the network, and mutation-level features to capture the potential of mutations to alter the architecture of the network. Though protein-level network features are shared by all variants in a protein, they nonetheless can interact with other amino acid level features to support classification; in all cases, combining both protein and mutation-level network features with classic amino acid features outperformed combining only mutation-level network features with classic amino acid features. Mutation-level features were helpful for distinguishing between variants within proteins, however because we designed them to capture the potential of mutations to alter the network architecture, all surface non-interface variants received the same value for these features.

Though network features show fairly different distributions for different classes of variants (Figure 4) and are orthogonal to the features typically used for variant classification, the best classifier combining both feature types show only modest gains over classifiers that use only classic amino acid features. This may result from more limited availability of training set mutations due to the requirement for structure and interface information to estimate network feature values. This requirement also constrained the coverage of benchmark set mutations that could be classified, though the values remained generally high, and over 70% in the worst case. Performance generally improved when we included predicted protein interactions from Interactome INSIDER [39], suggesting that *in silico* approaches may be an effective strategy to boost performance until more complete experimentally derived interaction maps are available.

Network features were more informative in the context of somatic mutations than when classifying inherited variants, though performance gains were observed in both cases. This is perhaps expected since inherited disease genes tended to be less central in PPI networks than cancer genes (Figure 2), and location at an interface by itself was less discriminatory in the germline setting (Figure 3D). We speculate that these differences may arise from different selective pressures acting on somatic versus germline variation. Because development at the organismal level is likely dependent on the integrity of molecular interaction networks, both pathogenic and neutral variants may be more constrained by network architecture; whereas in cancer, where selection operates at the cellular level and is predominantly positive, mutations may be better tolerated and be more advantageous in central network positions. We note also, however, that training sets for the driver versus passenger classification problems tend to be more gene-centric leading to concerns over whether cancer mutation classifiers distinguish primarily between genes [47]. Although we made an effort to include both drivers and passengers in each driver gene to mitigate this, it may still be reflected in the higher utility of protein-level network features for driver classification.

In conclusion, our study suggests that information about molecular interaction networks can be incorporated into machine-learning based variant interpretation frameworks. This opens future directions for the development of novel features capturing network information. Since networks can be constructed to model cell-type and condition specificity [48], it may be possible to build classifiers that can capture context-specific effects of variants. Furthermore, as studies have shown different interfaces are associated with different protein activities, network-based features could make it possible for machine-learning methods to provide more insight into the potential for mutations within a protein to have distinct functional consequences. We anticipate such advances will boost the utility of variant classification tools for precision medicine applications.

## MATERIALS AND METHODS

### Source of protein interaction data

To analyze disease gene centrality, we obtained a human PPI network of 12,811 proteins that are involved in 97,376 experimentally verified interactions with a confidence score higher than 0.4 from STRING v11.0 [29].

### Disease genes

A list of 125 high confidence cancer genes consisting of 54 oncogenes and 71 tumor suppressor genes was obtained from Vogelstein *et al*. [30]. We also obtained a list of 16,158 Mendelian genes from the OMIM database [49]. These genes were used to evaluate disease gene centrality (Figure 2).

### Collecting structural protein and protein interaction data

We obtained human protein interaction data (complete set) from Interactome3D [50], which contains a collection of a highly reliable set of experimentally identified human PPIs. We collected experimental co-crystal 3D structures for 5865 of these interactions from the Protein Data Bank (PDB) [51] and homology models for 5768 additional interactions [50] making a total of 11,633 interactions between 6807 proteins with structural protein and interaction data.

### Creating a structurally resolved PPI network

Amino acid residues were annotated as participating in a protein interaction interface based on KFC2 [52] scores, and we removed interactions containing fewer than 5 interface residues on either partner. Additionally, we calculated relative solvent accessible surface areas (RSA) using NACCESS [53] for all residues in each protein structure. Residues with RSA<5% and RSA>15% were designated as core and surface residues respectively. Residues with RSAs between these thresholds were excluded from further analysis due to ambiguity. When multiple PDB chains were available for the same protein, we used the consensus designation as the final label. The mapping of PDB residue positions onto UniProt residue positions was performed via PDBSWS web server [54]. After this mapping, we created a structurally resolved PPI network (named SRNet) of 6230 proteins and 10,615 protein-protein interactions with a total of 530,668 interface residues. To extend coverage of the structurally resolved network, we defined an extended network based on the “High Confidence” dataset of Interactome INSIDER [39], a human PPI network of 14,445 proteins with 110,206 interactions containing *in silico* interface residue predictions in addition to those derived from 3D structures.

### Source of somatic mutation data

To investigate structural location of cancer mutations on proteins (Figure 3), we mapped more than 1.4 million somatic missense mutations from TCGA [33] onto the structurally resolved PPI network using structural annotations. Only mutations mapping to canonical proteins were used. After this mapping, we identified a total of 56,667 interface residues of 5005 proteins that are involved in 9235 interactions as mutated.

### Training set for cancer mutation prediction

We collected a set of cancer missense mutations designated as likely-driver (n=2051) and likely-passenger (n=623,992) from Tokheim *et al*. [38]. Of these, 961 driver mutations from 32 genes and 28,043 passenger mutations from 2986 genes mapped to SRNet for a total of 29,004 mutations. All 32 genes with driver mutations also contain passenger mutations. To handle the driver vs. passenger mutation count imbalance in the training set by maintaining an approximate 1:4 driver vs. passenger mutation ratio similar to Carter *et al*. [42] while not overrepresenting particular genes, we limited the number of passenger mutations for each gene to 16 (median per gene driver mutation count) and collected 4613 passenger mutations at random across all genes with passenger mutations. In the extended network, we mapped 1513 driver mutations from 52 genes and 118,777 passenger mutations from 4478 genes for a total of 120,290 mutations. Thirty-eight of 52 genes with driver mutations also contain passenger mutations. To maintain an approximate 1:4 driver vs. passenger ratio as described above, we limited the number of passenger mutations for each gene to 17 (median per gene driver mutation count) and collected 6549 passenger mutations at random across all genes with passenger mutations.

### Training set for pathogenic variant prediction

We collected 5608 ‘pathogenic’ variants from ClinVar [34], and 4292 neutral variants including ‘common’ variants (allele frequency>1%) from EXAC [35], variants with ‘polymorphism’ classification from SwissVar [36] and ‘benign’ variants from ClinVar [34], that map to SRNet, totaling 9900 missense variants. We also collected 21,819 pathogenic, and 35,522 neutral variants from the same databases that map to the extended network, totaling 57,341 missense variants.

### Features

We designed 16 network-based features to quantify the potential impact of a mutation to the underlying network architecture, comprising 7 protein-level and 9 mutation-level features. The 7 protein-level features are degree, betweenness, closeness, eigenvector and load centralities, clustering coefficient, and pagerank of the proteins within the PPI network. They are computed using the NetworkX package of Python and they aim to characterize the centrality of a protein in the network based on measures such as the number of nodes it is directly connected to (degree), the amount of shortest paths it is involved in (betweenness and load), the overall closeness to all other nodes (closeness), its embeddedness (clustering coefficient), and the centrality of its neighbours (eigenvector and pagerank). 9 mutation-level features describe mutation 3D location (core, interface, surface) on the protein and changes in centrality of the protein within the PPI network resulting from mapping the mutation to network edges. Core mutations are assumed to affect all edges in the network, while interface mutations are mapped to corresponding edges in the network, and surface mutations retain all edges. The remaining 8 mutation-level features are based on this description of how the mutation perturbs the network by capturing degree change, betweenness change, closeness change, eigenvector change, clustering coefficient change, load change, pagerank change and percent degree change. Non-network related amino acid based features (n=83) obtained from the SNVBox database [37] describe substitution effects on amino acid biophysical properties, evolutionary conservation of variant sites, local sequence biases, and site-specific functional annotations. The number of PubMed studies featuring each gene was obtained from the NCBI database (https://ftp.ncbi.nih.gov/gene/DATA/gene2pubmed.gz) for all genes in SRNet with NCBI (Entrez) gene IDs. This was used to assess the potential for gene-level features to be affected by study bias.

### Classifier training

We trained a Random Forest classifier on the training set using the scikit-learn Python package. We used a forest with 1000 trees and default parameters. To avoid classifier overfitting, we performed prediction using a 5-fold gene hold out cross-validation by dividing the training set into 5 random folds for cross-validation while ensuring a balanced number of disease and neutral mutations across the folds. All mutations occurring in the same gene were kept within the same fold. The classifier score represents the percentage of decision trees that classify a mutation as a disease mutation (driver or pathogenic). Receiver Operator Characteristic (ROC) curves were constructed from the classifier scores and the AUC statistic was used as a measure of classifier performance. To compare the performance of different features for identifying disease mutations, we trained different classifiers on different sets of features: all 16 network-based features (Net), dividing network-based features into 7 protein-level features (Prot) and 9 mutation-level features (Mut), 83 non-network amino acid (AA) features, 83 amino acid features combined with 9 mutation-level features (AA & Mut) or 83 amino acid features combined with all 16 network features (AA & Net). Pearson correlation coefficient was used to evaluate feature correlations.

### Benchmark datasets

We obtained 4 pan-cancer benchmark sets of missense mutations consisting of an *in vivo* screen: Kim *et al*. [40], an *in vitro* assay: Ng *et al*. [41], and 2 literature-derived datasets: MSK-IMPACT and CGC-recurrent from Tokheim *et al*. [38]. The *in vivo* screen contains 71 mutations selected based on their presence in sequenced human tumors and screened in mice to assess oncogenicity and then labeled as ‘functional’ or ‘neutral’ based on their abundance [40]. The *in vitro* assay consists of 747 mutations from a growth factor dependent cell viability assay annotated as ‘activating’ for increased cell viability, or as ‘neutral’ for the remaining, with the assumption that a mutation yielding higher cell viability indicates driverness [41]. The MSK-IMPACT dataset is composed of mutations from approximately 10,000 tumors [55] on 414 cancer-related genes (MSK-IMPACT gene panel) labeled as positive class if annotated as ‘oncogenic’ or ‘likely oncogenic’ in OncoKB [56], or as negative class if not. The CGC-recurrent dataset consists of TCGA mutations annotated as positive class if recurrent in a set of curated likely driver genes from the Cancer Gene Census [57], or as negative class if not.

## Comparison to other methods

Performance was compared to 8 state-of-the-art methods that do not use network-based information. Predictions for 4 cancer-focused methods: CHASM [42], TransFIC [43], FATHMM [44], and CanDrA [45], and 4 population-based methods: SIFT [4], Polyphen2 [5], VEST [46], REVEL [8], on benchmark sets were obtained from Tokheim *et al*. [38]. Classifier performance was compared using AUC statistics from both ROC and Precision-Recall (PR) curves.

## Acknowledgements

This work was supported by SDCSB/CCMI Systems Biology training grant (GM085764 and CA209891) to K.O. and NIH grant DP5 OD017937 and CIFAR award FL-000655 to H.C. NIH grant 2P41GM103504-11 provided access to computational resources. The results herein are in part based upon data generated by the **TCGA** Research Network: https://www.cancer.gov/**tcga**.

## Supporting Information

**Figure S1. Study bias analysis**. Correlation of node degree (the number of interacting partners) of proteins in SRNet with the number of PubMed publications they appear in (Pearson r=0.23, p-value=4.78e-54).

