## Supplementary material for "Predicting functional consequences of mutations using molecular interaction network features": Figure S1

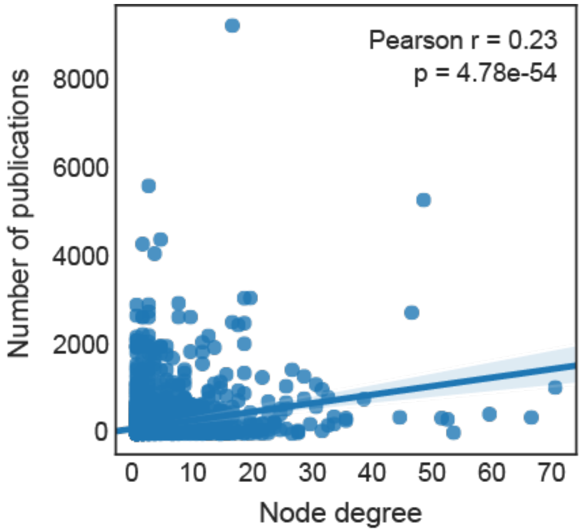


**Figure S1. Study bias analysis.** Correlation of node degree (the number of interacting partners) of proteins in SRNet with the number of PubMed publications they appear in (Pearson r=0.23, p-value=4.78e-54).
